# BowSaw: inferring higher-order trait interactions associated with complex biological phenotypes

**DOI:** 10.1101/839357

**Authors:** Demetrius DiMucci, Mark Kon, Daniel Segrè

## Abstract

Machine learning is helping the interpretation of biological complexity by enabling the inference and classification of cellular, organismal and ecological phenotypes based on large datasets, e.g. from genomic, transcriptomic and metagenomic analyses. A number of available algorithms can help search these datasets to uncover patterns associated with specific traits, including disease-related attributes. While, in many instances, treating an algorithm as a black box is sufficient, it is interesting to pursue an enhanced understanding of how system variables end up contributing to a specific output, as an avenue towards new mechanistic insight. Here we address this challenge through a suite of algorithms, named BowSaw, which takes advantage of the structure of a trained random forest algorithm to identify combinations of variables (“rules”) frequently used for classification. We first apply BowSaw to a simulated dataset, and show that the algorithm can accurately recover the sets of variables used to generate the phenotypes through complex Boolean rules, even under challenging noise levels. We next apply our method to data from the integrative Human Microbiome Project and find previously unreported high-order combinations of microbial taxa putatively associated with Crohn’s disease. By leveraging the structure of trees within a random forest, BowSaw provides a new way of using decision trees to generate testable biological hypotheses.

## Introduction

The production of large biological data sets with high-throughput techniques has increased the utilization of supervised machine learning algorithms to produce predictions of complex phenotypes (e.g. healthy vs. disease) from measurable traits. These algorithms use measurements of relevant traits such as gene variants, the presence/absence of microbial taxa, or metabolic consumption variables as predictors. Categorical prediction of phenotypes is typically the end goal of these applications. However, an additional benefit of these algorithms is the potential to extract explanatory classification rules. In this context, a rule is defined as a Boolean function of a set of traits, such that the value of the function is 1 (true) when the traits are associated with a given phenotype. Identifying the relationships between the traits involved in classification rules may yield key insights into the biological processes associated with important phenotypes [1, 2]. This realization is creating demand for methods that assist in the interpretation of supervised machine learning methods [3–5], especially when the measured traits may be causal agents of disease states, such as genetic variants or microbial taxa [6]. Identifying classification rules associated with a phenotype of interest is valuable because these rules are likely to carry information about the causal mechanisms that generate the phenotype.

Algorithms that are particularly valuable in this respect are those involving decision trees, such as random forests, since decision trees are easily interpretable [7]. Decision trees are rule-based classifiers, where rules arise from a series of “yes-no” questions that can efficiently divide the data into categorical groups. In a biological context, such rules may arise from sets of genes whose simultaneous modulation could affect a phenotype, or sets of microbial species whose co-occurrence may be associated with a disease state. While in several cases it seems like disease phenotypes are uniquely associated with a single specific pattern (e.g. retinoblastoma [8]), there is increasing evidence for cases in which multiple distinct patterns can be associated with (and potentially causing) the same high-level phenotype [9, 10]. A particular example we will explore in this work is the multiplicity of distinct microbial presence/absence patterns which may be associated with Crohn’s disease [11]. Crohn’s disease has five clinically defined sub-types [12] but studies of the associated microbiome do not usually indicate which form of Crohn’s disease a donor has been diagnosed with. Each sub-type of the disease may be associated with different microbes, each requiring different treatment regimes. Thus, identifying rules associated with sub-populations within a given phenotype label are of great interest due to potential therapeutic implications.

The fact that there may be multiple etiologies that generate the same or similar phenotypes complicates the straightforward interpretation of parameter coefficients or variable importance scores [13, 14]. Uncovering the multiple interactions between predictive variables as they relate to phenotypic labels remains a challenging statistical endeavor, but one that is of paramount importance. Identifying the associated rules that a random forest uses to classify a given sample as having a particular disease enables the development of mechanistic hypotheses for follow up-studies. This challenge, and an overview of the key strategy we propose, are illustrated in Figure 1. In figure 1A we depict a toy model where measured variables (traits) have only two possible values (e.g.: present/absent), the high-level phenotype (category) is binary (e.g.: no disease/disease), and two distinct Boolean rules can both generate the phenotype. The goal in this case is to identify each of the rules that are associated with the phenotype. The multiple Boolean rules obtained in this manner can be thought of as a consensus decision tree that possesses the most informative branches of the forest with respect to a given class label. In this work, we will show how this can be achieved by in-depth analyses of any given random forest (RF) (Fig. 1B).

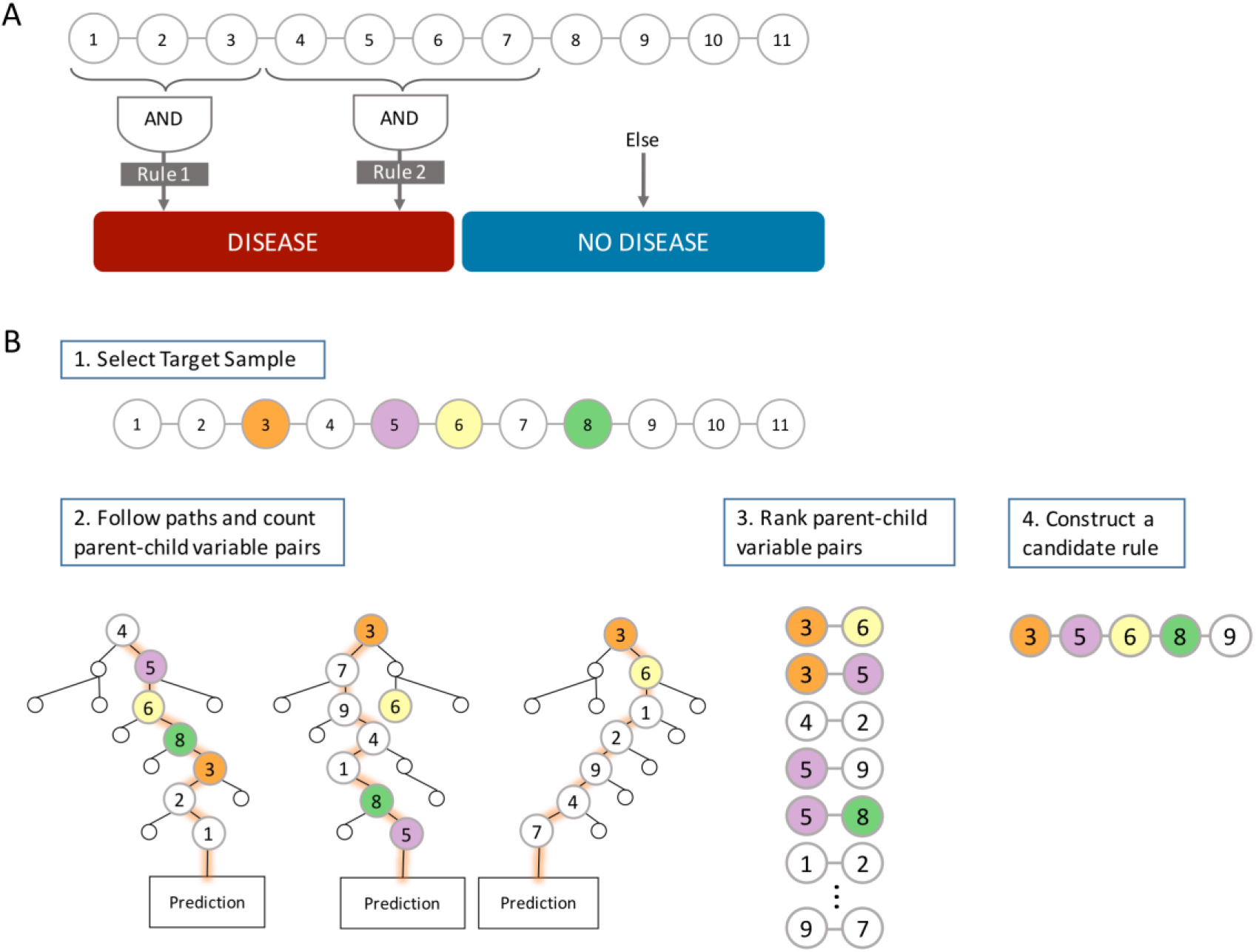
**A** In a hypothetical dataset there may be two phenotype labels – “Disease” and “No Disease”, that we wish to discriminate based on input predictor variables. In this example, there are two distinct high-order patterns that both confer the “Disease” phenotype. Our goal is to identify a potentially diverse set of patterns (or, in this simplified case, all patterns) that are associated with the “Disease” label. **B** Conceptual pipeline of BowSaw. In (1) we begin by identifying the vector of a target instance that has the target observed label. In this example, the colored nodes indicate a true associated pattern, which is unknown to us. In (2) we follow the path of the instance through each of its out-of-bag trees and record how often the sample encounters sequential pairs of variables. (3) Each ordered pair sequence is sorted in descending order by its observed frequency. (4) Starting from the top of the list, pair sequences are iteratively evaluated and added to an undirected network of variables (i.e. a candidate rule) until this network is maximally associated with the observed phenotype of the target vector or the list of ordered pairs is exhausted. Each sample with the label of interest yields one such candidate rule. These rules are then aggregated and curated to obtain a concise set of rules that explain class-specific classification decisions that occur in the forest.

The random forest algorithm intrinsically takes advantage of non-linear relationships between variables and is widely used in the life sciences [15–17]. RFs, when used to distinguish between disease states known to have multiple causes, often result in excellent classifiers [18, 19]. It has also been reported that RFs capture subtle statistical interactions between variables [13]. Unfortunately, an RF is not straightforwardly interpretable despite its hierarchical structure, and recovering those interactions is notoriously difficult [14] due in large part to the method’s reliance on ensembles of trees [20]. The difficulties in interpretation created by these properties has led many to refer to RF as a ‘black box’ model [21].

Identifying the rules that a RF utilizes in classification tasks is an active area of research, and many strategies have been developed to address this problem. Effective strategies have focused on evaluating how individual variables influence the classification probabilities of specific samples [22, 23], pruning existing decision rules found in the tree ensemble to produce compact models [24], computing conditional importance scores [25], or iteratively enriching the most prevalent variable co-occurrences through regularization [26]. These approaches offer valuable methods for the identification of statistical interactions between variables. However, we and others have observed that while these methods are capable of recovering a true causal rule in simulated data when exactly one such rule is present, the existence of multiple rules associated with one phenotype can confound interpretation efforts [26].

Here we describe BowSaw, a new set of algorithms that utilizes variable interactions in a trained RF model in order to extract multiple candidate explanatory rules. With BowSaw, we set out to develop a *post hoc* method intended to aid in the discovery of these rules when the input variables are categorical in nature. The primary approach of BowSaw is to start by approximating a best combination of variables (i.e. a rule) that explain the forest’s predictions for individual instances of a given class in the data set and then to curate the collection of best combinations to obtain a concise set of combinations that collectively segregate a class of interest with high precision. For individual instances a rule is identified by systematically quantifying the co-occurrence of specific variable pairs across trees in the forest that attempt to predict the class of the instance (out-of-bag trees) and then using the frequency of co-occurring variable pairs to guide the construction of a rule that precisely identifies the instance as its observed class. For the entire set of instances, we then curate the collection of all rules identified this way in order to produce a small set of rules that are broadly and precisely applicable to instances of the given class label.

We first demonstrate that BowSaw can recover true rules by applying the algorithms to simulated data sets of varying complexity. We then apply BowSaw to a study on the role of the gut microbiome on Crohn’s disease [11], and show that it can find a previously unreported combination of microbial taxa that is broadly and precisely associated with Crohn’s disease instances in the data set. In its current implementation BowSaw can be applied to any dataset with categorical or discrete predictors with any number of class labels.

## Methods

### Overview of the pipeline

Provided with a trained random forest and a training set, BowSaw goes through three steps in order to generate a candidate rule (variable-value combination) for each observation associated with the phenotype of interest. First, for a specific observation, the *Count* algorithm counts the frequency of unique ordered pairs of variables encountered along each of its out-of-bag trees in the forest (Figure 1B – step 2). Second, for that observation, the *Construct* algorithm takes the counts from the first step and generates a list of ordered pairs, ranked by their frequencies, then uses this list as a guide to construct a candidate decision rule (which could consist of two or more variables) that is maximally associated with the observed phenotype (Figure 1B – steps 3 – 4). Finally, the *Curate* algorithm pools the candidate decision rules from each observation together in order to select a subset of rules that collectively account for all of the samples with the desired phenotype (Figure 1B – step 5). Optionally, the *Sub-rule* algorithm can be used to generate pruned versions of candidate rules prior to applying the Curate algorithm in order to obtain a more concise, albeit less specific, set of candidate rules. The Count and Curate algorithms generate the candidate rules for individual observations while the Curate and Sub-rule algorithms produce a combined set of rules that account for all observations with the chosen phenotype.

In the following section, we provide a description of the inputs BowSaw takes and the algorithms that implement these steps along with pseudocode.

### Inputs

BowSaw takes as inputs a dataset, ***D***, composed of *N* observed vectors ***x***_*i*_ (together with their respective classes *k*_*i*_) each of *p* categorical variables. There are assumed to be *K* possible class labels for each vector in ***D*** which for the purposes of this discussion denote different phenotypes. A random forest is assumed to be trained on ***D*** to distinguish the classes *k* = 1, …, *K*. Additionally, BowSaw takes as input the feature vector ***x***_*i*_ of a specific observation for which the goal is to identify a set of simplified rules associated with the phenotype *k*_*i*_.

### Counting stubs

Given an RF machine ***M*** trained on dataset ***D*** and a feature vector *x* = (*x*_1_, *x*_2_, …, *x*_*p*_) ∈ ***D***, the first sub-routine of our method (the *count algorithm*) proceeds as follows. It starts by identifying among the set of trees in ***M***, those sub-paths (sequences of successive variable indices) encountered by sample ***x*** as it travels through ***M***_*x*_, its set of out-of-bag trees. An out-of-bag tree is a tree for which *x* was not included in the training set. For a specific path ***P*** in ***M***_*x*_ the sequence of successive variable indices forms a vector ***ν*** = (*ν*_1_, …, *ν*_*r*_) (note that each *ν*_*j*_ is one of the variables *x*_*j*_). Each stub (ordered pair of sequentially encountered variables *ν*_*i*_*ν*_*i*+1_) in all out-of-bag along ***p*** for *i* = 1, … *r*<u>−1</u> is accounted for in a *p* × *p* matrix ***C***^***x***^, where the element 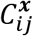 records the number of stubs containing the ordered pair of variables *x*_*i*_ and *x*_*j*_ among all paths of ***M***_*x*_.

**Algorithm 1:**
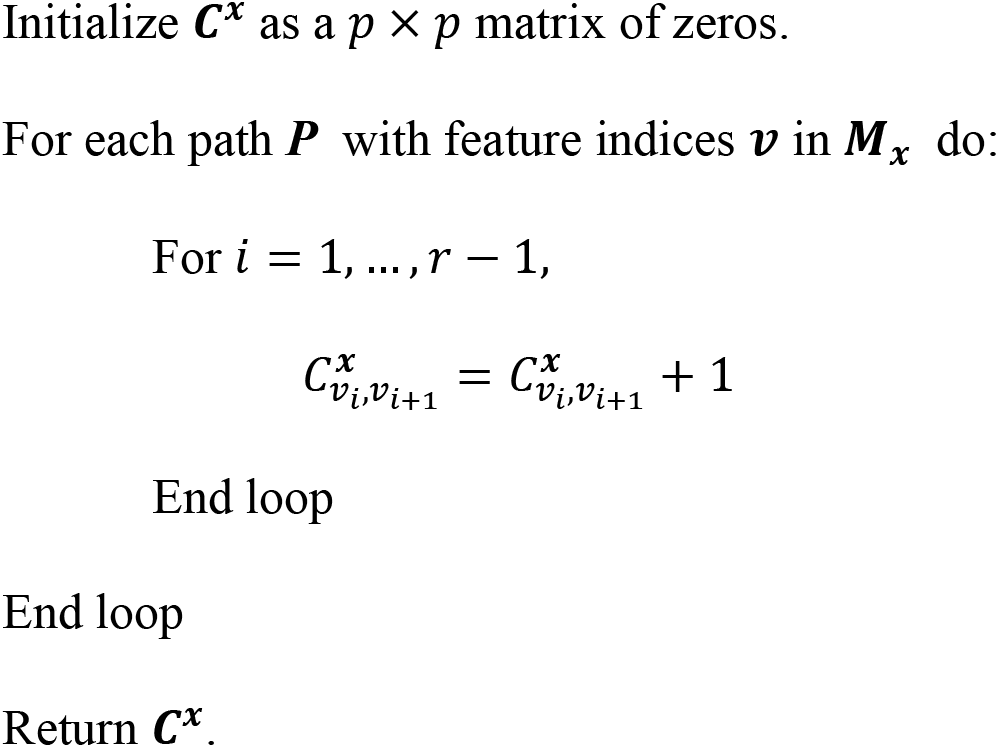
*Count Algorithm* Pseudocode.

For simplicity, henceforth we will denote ***C*** = ***C***^***x***^, remembering that ***C*** continues to depend on the fixed sample ***x***.

### Constructing a candidate rule

A *rule* for classifying to a test point ***x*** will have the form “***x***_***I***_ = ***a***_***I***_ implies ***x*** is in class *k*”. Here ***I*** is a designated subcollection of the variable indices *i* = 1, …, *p*, and 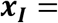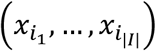 is the sub-vector of current vector ***x*** = (*x*_1_, …, *x*_*p*_) corresponding just to the indices *i*_*j*_ ∈ ***I***. The vector 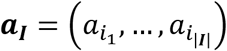 will denote an assigned set of values to the *x*_*i*_, i.e., so that *x*_*i*_ = *a*_*i*_ for *i* ∈ ***I***. Thus the condition ***x***_***I***_ = ***a***_***I***_ means assignment of values to *x*_*i*_ for *i* ∈ ***I***. The rule is that if training vector ***x*** satisfies ***x***_***I***_ = ***a***_***I***_, we classify ***x*** into category *k*.

The second sub-routine (the *construct algorithm*) builds a candidate rule ***R***, based (initially) on a fixed training point, say ***a*** ∈ ***D***, in class *k*. This is done by first placing all of the stubs (*i*, *j*) with non-zero counts ***C***_*ij*_ into a list ***L*** sorted in descending order by their values in ***C***.

We define the candidate rule ***R*** (based on ***a***) through the following steps. We initialize using the first stub *L*_1_ = (*i*_1_, *j*_1_) in the list ***L***, together with the two fixed values 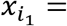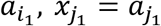. This is the initialized form of the rule ***R***, which requires that for any test vector, its values at the above indices *i*_1_ and *j*_1_ match the values of the above fixed training vector ***a*** ∈ ***D***, so that 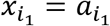, and 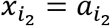. For brevity, denote the pair (*i*_1_, *j*_1_) = *I*_1_ and the corresponding assigned values as 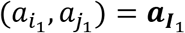. Then the content of rule ***R*** will be denoted succinctly as ***R***: ***x***_***I***_ = ***a***_***I***_ ⇒ class *k*. Since ordering of the indices *i*_1_, *j*_1_ does not matter, (as long as the indices are identified), we will henceforth write (*i*_1_, *j*_2_) → {*i*_1_, *j*_2_}.

We then update rule ***R*** as follows. We find all ***x*** ∈ ***D*** that satisfy the initial part of rule ***R***, i.e., ***x***_***I***_ = ***a***_***I***_ i.e., all training points matching the two indices {*i*_1_, *j*_1_} of training sample ***a***, and store them as a subcollection ***D***_1_ ⊂ ***D*** of the training set. We call *F* the fraction of data points in ***D***_1_ that have phenotype *k*, i.e., match the phenotype of the initial sample ***a*** ∈ ***D***. If ***F*** = 1, we stop and return the current above rule ***R***. If ***F*** < 1, we continue by choosing the second stub *L*_2_ = {*i*_2_, *j*_2_} in the above list ***L***, and augment the current rule ***R*** by adding the condition 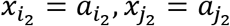 (again written 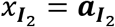) and maintaining the assignment of class *k* (i.e., the same class as the currently fixed sample ***a*** ∈ ***D***). If the second stub *L*_2_ happens to overlap with the initial stub *L*_1_, this added condition in the rule ***R*** will clearly be consistent, being still based on the fixed sample ***a***. We augment the current index list ***I***_1_ to a l ist ***I***_2_, adding to it the two new indices *i*_2_ and *j*_2_, so that now ***I***_2_ = {*i*_1_, *j*_1_, *i*_2_, *j*_2_} writing the augmented rule as 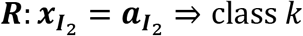. Again defining *F* to be the fraction of the data subset ***D***_2_ (matching the more restrictive new rule ***R***) with phenotype *k*, we stop the algorithm and use the current rule ***R*** if *F* = 1, and otherwise augment rule ***R*** by adding the indices *L*_3_ = (*i*_3_, *j*_3_) to it, as above, yielding a larger set ***I***_3_ of indices and the augmented rule 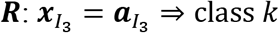, with a more restricted subset ***D***_3_ ⊂ ***D***, and a new value for *F*, now the fraction of ***D***_3_ in the class *k* of the fixed ***a*** ∈ ***D***.

This process continues until the fraction *F* = 1, i.e., 100% of the samples in ***D*** match the current set of indices, and also match the class ***a*** of the current sample ***a***. Alternatively, the algorithm stops when all stubs in ***L*** have been exhausted.

**Algorithm 2:**
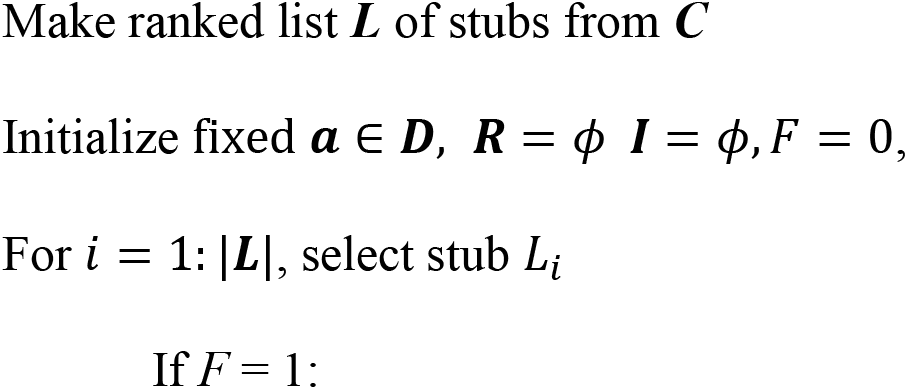
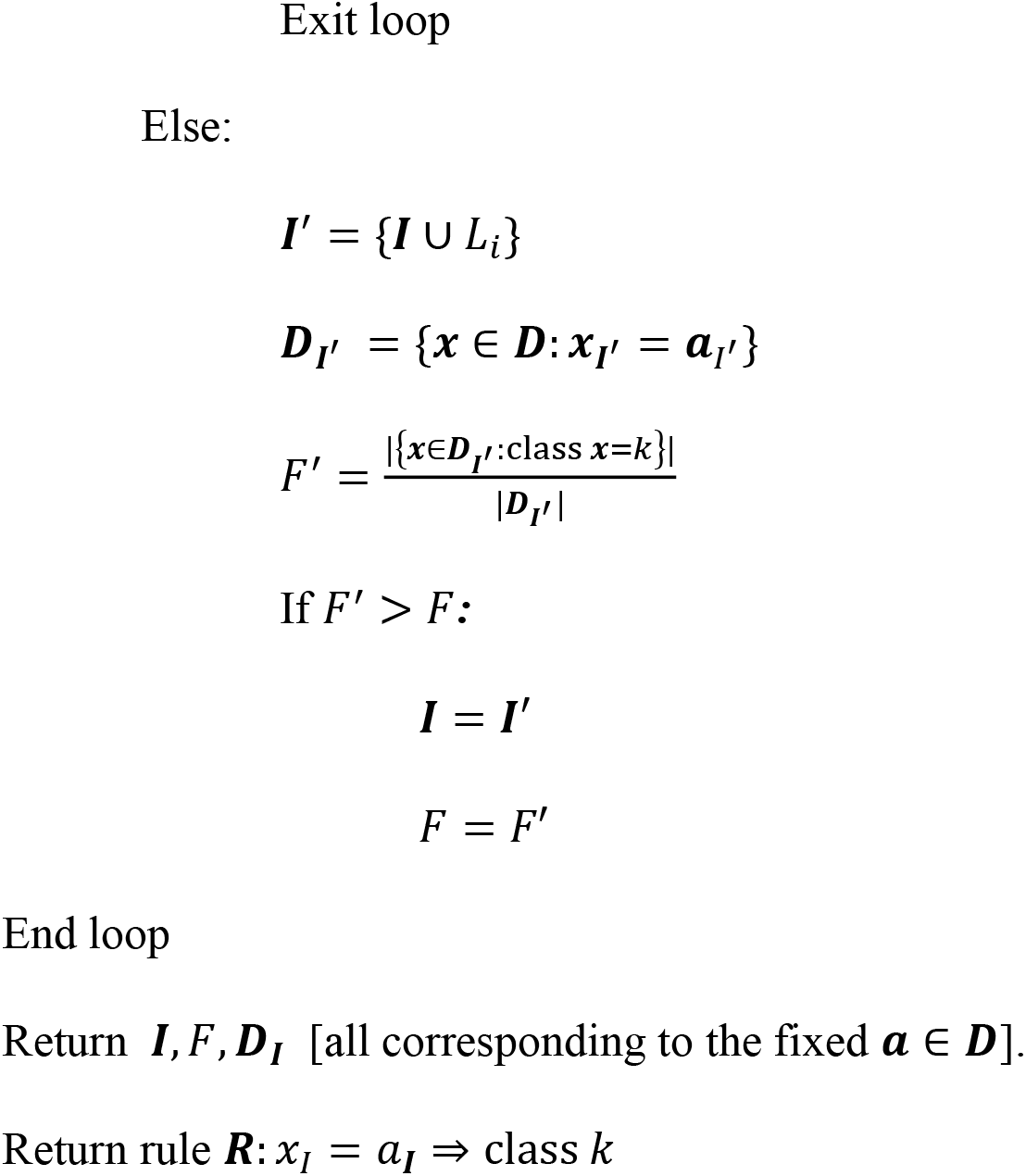
*Construct Algorithm* Pseudocode.

### Curating candidate rules

The *count* and *construct* algorithms are the heart of BowSaw. In our workflow, we apply these algorithms to each observation ***a*** ∈ ***D*** that has the desired observed phenotype *k*. We call the set of these vectors ***D***^*k*^ ⊂ ***D***. By default, we produce a single candidate rule for each vector in ***a*** ∈ ***D***^*k*^. We store each candidate rule in list ***Q*** and rank them by their respective values of |***I***|, i.e., the number of indices in the respective rules. Since ***Q*** may include many redundant rules, we developed another sub-routine (the *curate algorithm*) to generate a concise set of candidate rules that collectively account for all data vectors ***D***^*k*^ in class *k*. Briefly, we initialize an empty list ***E***, to which we add the top ranked rule from ***Q*** (by default this is the rule with the greatest value of |***I***|), and record the index of samples in ***D*** that match any rule in ***E*** and also have the desired observed phenotype class *k*, into a set ***A***. Next, we determine how many samples remain unaccounted for, i.e. are in ***U*** = ***D***^*k*^~***A***, Then we determine which of the remaining rules in ***Q*** minimizes |***U***|, add it to ***E***, and repeat these steps until ***U*** is an empty set.

**Algorithm 3:**
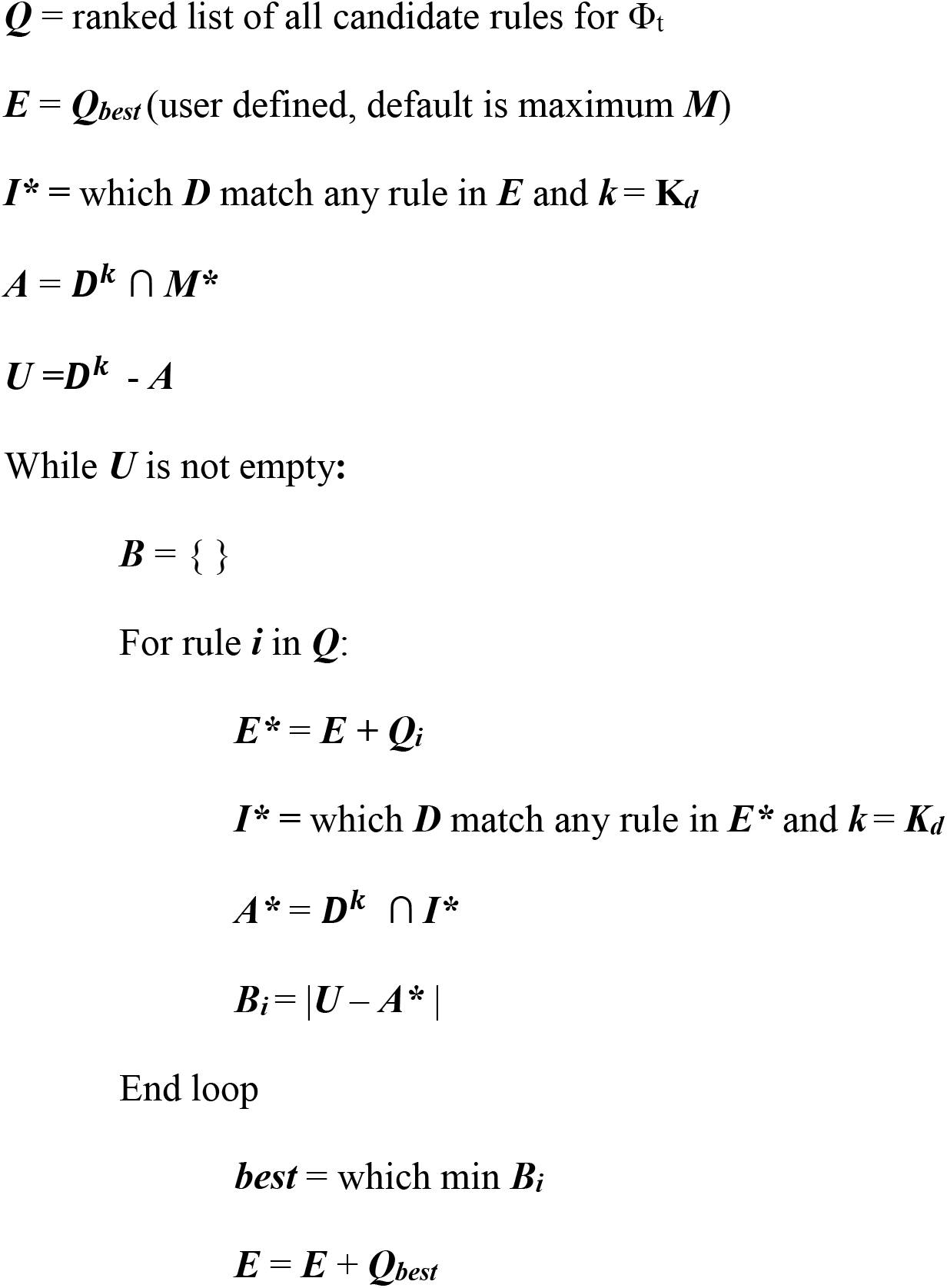
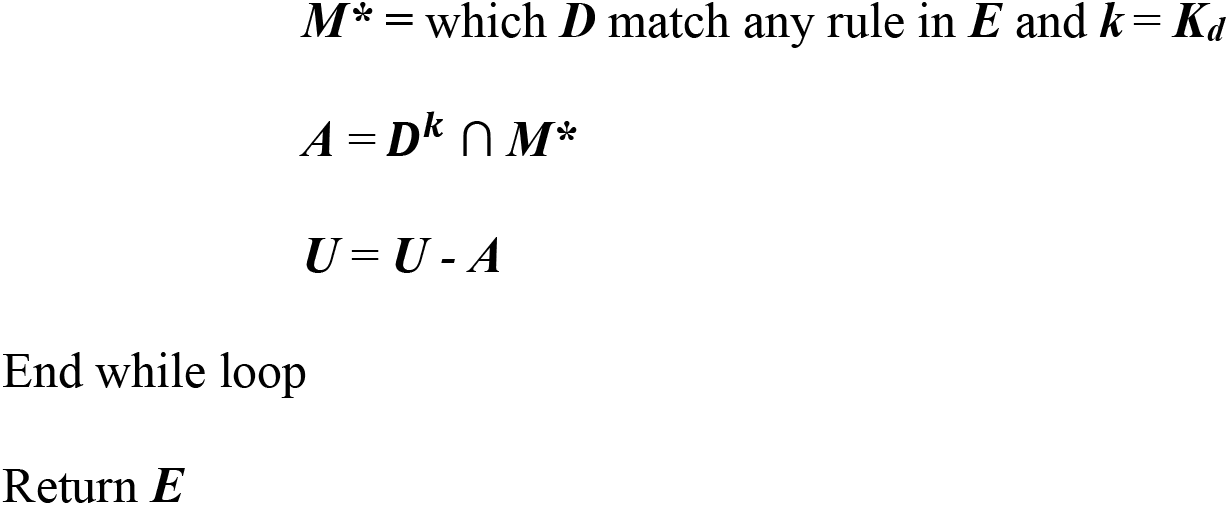
*Curate algorithm* pseudocode.

### Constructing sub-rules

Since rules are rarely 100% associated with any given phenotype, we devised a strategy for selecting a set of candidate sub-rules that account for all samples with desired observed phenotype class *k*. Candidate sub-rules are shorter candidate rules derived from larger candidate rules by omitting one or more variables. For each candidate rule in ***E***, we identify sub-rules that meet a user-defined complexity criteria, e.g. only produce sub-rules that are composed of three or four variables and their corresponding values. We place each of the unique sub-rules into a new list ***E***_*sub*_. Then the corresponding number of identical matches, ***I***, and proportion of ***I*** that have the phenotype ***K***_*d*_, ***F***, are determined. At this stage, we can apply our third sub-routine (the *Curate* algorithm) to ***E***_*sub*_ to obtain a parsimonious list of sub-rules that accounts for **x**_**all**_. In our pipeline, we also choose thresholds based on desired levels of ***I*** and/or ***F*** in order to eliminate poor candidate sub-rules from consideration. In this study, we decided on the thresholds after visually inspecting a plot of ***F*** against ***I***.

**Algorithm 4:**
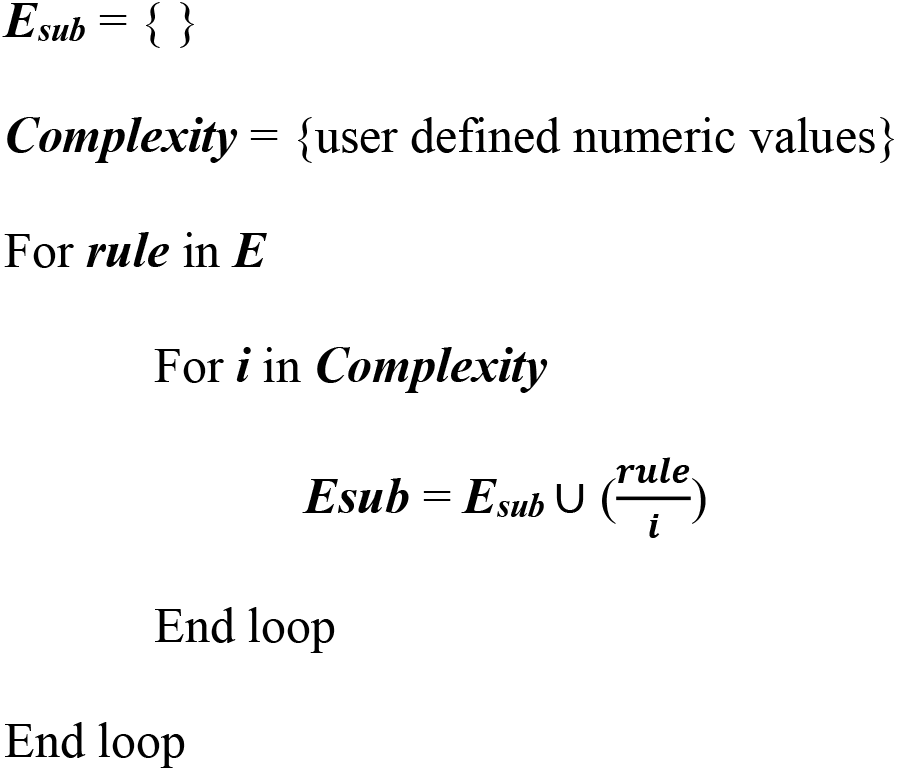
*Sub-rule algorithm* pseudocode.

The algorithms described above are generalizable to multi-classification tasks but are currently limited to discretized or categorical representations of the feature space. Pseudocode for implementing each of the algorithms described above along with an implementation of the algorithms in R [27] can be found in the supplemental files and on github: https://github.com/ddimucci/BowSaw.

## Results

### Application to simulated Data

To test the capacity of BowSaw to recover multiple decision rules, we applied it to increasingly challenging simulated data sets. These data set consists of binary vectors representing different observations. The phenotype associated with each observation is a function of the corresponding vector. The function consists of a set of multiple mutually distinct Boolean rules, such that if a rule is satisfied, it will cause the observation to have the phenotype with a certain probability (which we call here “penetrance” because of its resemblance to the genetics concept). The first dataset (IDEALIZED) we use is relatively simple, and includes multiple equally prevalent rules. It is also generated under the assumption that there are no unmeasured confounders, i.e. that if an observation does have a phenotype, then it must be satisfying at least one of the above rules. We then apply BowSaw to a more challenging scenario (INTERMEDIATE) in which the phenotype-generating rules differ in their relative prevalence and the assumption of unmeasured confounders is violated. Finally, is a set of data sets with complex co-varying parameters (COMPLEX), we systematically varied the underlying parameters of the simulation and examined the relationship between summary statistics of the RF performance and the ability of BowSaw to generate candidate rules containing the true phenotype-generating rules.

For the IDEALIZED scenario, we simulated data set of 100 independent and identically distributed random binary variables and 2,000 observations. We randomly defined five rules that each required four randomly selected variables each to have specific values (e.g. all variables equal to 1) in order to assign a hypothetical phenotype with likelihood between .8 and .9. Here we present the results of this scenario with a specified random seed, but other seeds and parameters can be explored using the scripts provided in the supplemental files. Using these parameters 479 samples were assigned the phenotype and BowSaw produced a set of 135 unique candidate rules ranging in complexity from six to fourteen variables. From these rules, we produced all sub-rules ranging involving anywhere from two to five variables, which resulted in unique 50,034 sub-rules. We calculated the number of matches |***I***|, the proportion of samples with the phenotype, ***F***, for each sub-rule, and visualized these values in order to select an association threshold (Figure 2A). To reduce the number of sub-rules that the curate algorithm would need to examine, we eliminated from consideration any rules that had an ***F*** below 80%. We selected an 80% threshold because in the cluster centered around 125 matching samples there is a small cloud of rules that are clearly segregating the phenotype more efficiently than the others are. We selected the sub-rule with largest |***I***| among these as the top candidate rule. This produced a final list consisting of five candidate rules that accounted for all of the samples with the phenotype and were each one of the true phenotype generating rules (Figure 3A red points). These results demonstrate that in an ideal scenario with no phenotype diagnosis errors, BowSaw is indeed capable of recovering multiple true rules.

**Figure 2.**
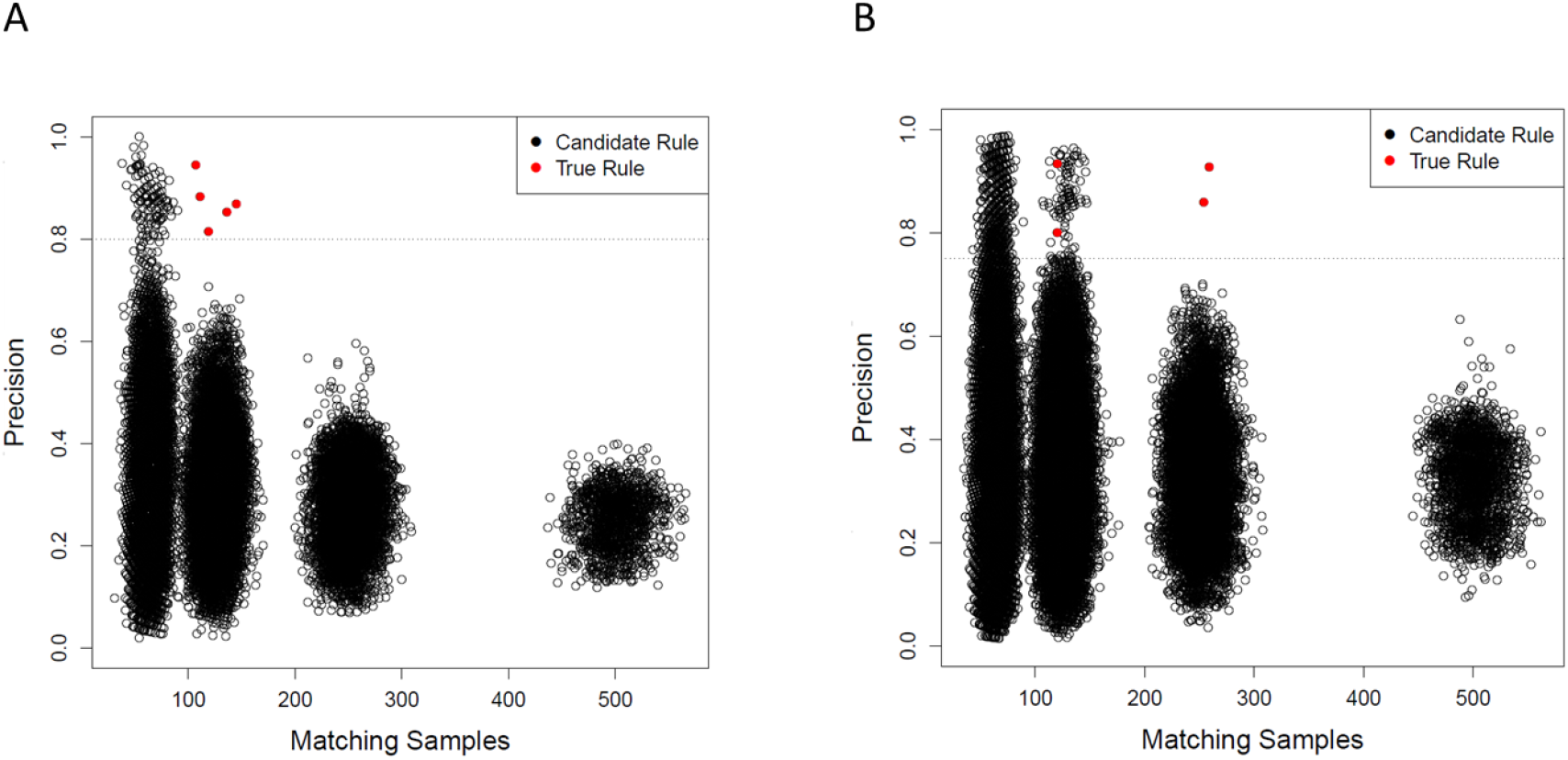
**A** Precision of candidate sub-rules against the number of exactly matching samples for the ideal scenario. Each point represents a unique sub-rule. X-axis is the number of samples that exactly match the pattern defined by the rule. Y-axis is the fraction of matching samples with the observed phenotype (i.e. precision of the rule). Each cluster of points corresponds to decreasing rule complexity from 5 variables per rule to 2 on the right most cluster. These clusters appear because the values of each variable is produced by an identical binomial distribution. Dashed line is the precision threshold we set. Only candidate rules with precision above this threshold were considered for the curate algorithm. Red points are the causative sub-rules we defined. BowSaw correctly identified all five red points in this scenario. **B** Candidate sub-rules generated for the more challenging scenario. We defined 5 causative rules of varying lengths in this scenario and allowed 2% of samples without a causative rule to be assigned the label. BowSaw completely 4 of the causative rules (red points). The longest rule which involved 5 variables was not recovered.

**Figure 3.**
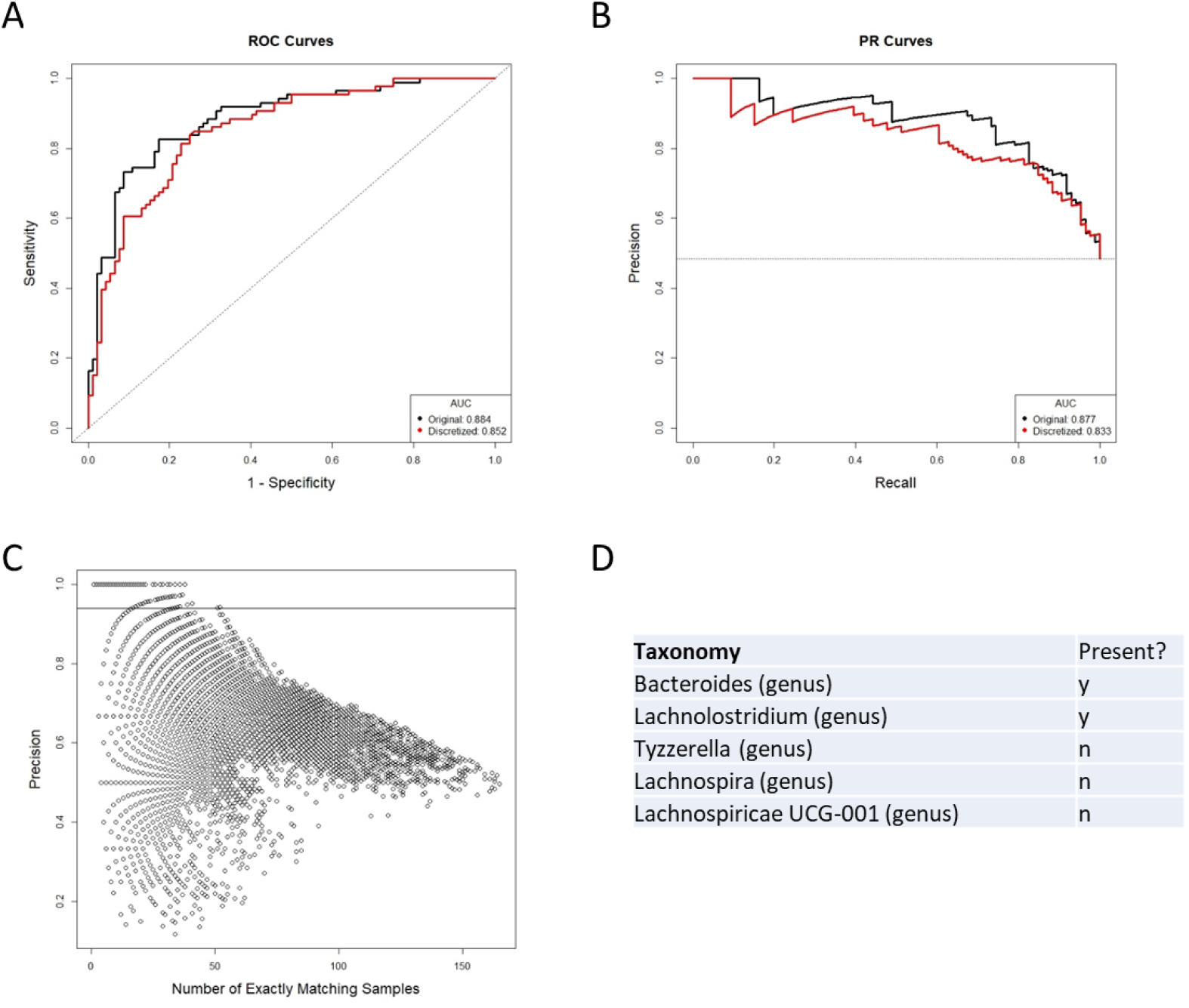
**A** Performance of the random forest classifier as measured by area under the receiver operator curve (ROC-AUC) is not strongly perturbed by simplifying OTU representation to a presence/absence scheme versus the original continuous count. Dashed line indicates the performance of a perfectly random classifier. **B** The area under the curve of the precision recall curve is similarly not strongly affected by the new representation scheme. Dashed horizontal line is the random performance line. **C** Each point represents a unique candidate sub-rule. On the x-axis is the number of samples in the data matrix that are subject to that rule. The y-axis represents what fraction of matching samples were diagnosed as Crohn’s disease. **D** The taxon identities of the OTUs that make up the most generally applicable of the sub-rules where all matching samples have the Crohn’s disease label.

For the more challenging scenario (INTERMEDIATE), we generated the data set the same as before except this time we allowed the five underlying rules to vary in complexity from three to five variables. Varying the complexities of rules resulted in different prevalence among them, as rules that are more complicated are less likely to appear in the data. In this case, we had one rule of complexity five, two that required four variables, and two that used three variables. We also added background noise by randomly assigning the phenotype to 2% of samples that did not possess any of the rules. BowSaw produced 176 unique candidate rules involving between six to thirteen variables. From this list we generated 68,938 sub-rules and chose an association threshold of 75% because there are two clusters at ~|***I***| = 125 that begin to clearly separate in that range and the two outlier points at ~|***I***| = 250 do not combine to account for all of the phenotype (Figure 3B). Applying the curate algorithm to the rules meeting this threshold produced 20 candidate sub-rules the top four (when ranked by |***I***|) of which were true rules. The rule of five variables was not recovered. These results show that BowSaw is able to recover strongly associated patters (and in this case, causal patterns) even in the presence of noise, but low prevalence rules can be masked by high prevalence rules.

We used the same data generation method to investigate BowSaw’s ability to produce candidate rules containing true rules when the underlying parameters change. We applied BowSaw to 20,000 simulated data sets where we randomly altered the number of features, sample size (200 or 2,000 samples), complexity of the rules, number of rules, the likelihood of each rule assigning the phenotype, and the background noise. We identified scenarios where rule recovery with BowSaw performs very well and situations in which it fails to recover any rules at all. Additionally, we found a strong linear relationship between BowSaw’s performance measured as the average fraction of rules recovered and the of number of samples, number of features, and two evaluation metrics for RF model – the area under the curve for both the receiver operator characteristic and precision recall curves (Figure S1).

### Application to Human Microbiome Data

Irregular distributions of microbial taxa within the gut are often associated with serious illnesses such as Crohn’s disease or ulcerative colitis [28, 29]. Human microbiome studies regularly use 16s sequencing methods and extensive reference databases to report on microbial taxa found in samples as operational taxon units (OTUs). RF classifiers are frequently built using counts of OTUs to accurately discriminate between disease and healthy patient samples [30, 31]. Despite their demonstrated effectiveness as good classifiers of Crohn’s disease, studies that look to discover associations with disease status typically focus on individual OTUs while specific microbial association rules found by RF are not discussed, as a result it is uncertain how heterogeneous study cohorts are. To investigate potential rule heterogeneity in a human microbiome cohort we downloaded processed files from the Human Microbiome Project for inflammatory bowel disease (IBD) [11] which contain information on the taxonomic profiles of 982 OTUs in 178 patients – 86 of which have been diagnosed with Crohn’s disease, 46 diagnosed with ulcerative colitis, and 46 diagnosed as non-IBD. We were specifically interested in finding rules that separate the Crohn’s disease samples from ulcerative colitis and non-IBD, so we framed the problem as a binary classification task with Crohn’s disease as the target phenotype.

Since the current implementation of BowSaw is limited to finding rules when the variables have categorical values, we first converted the OTU counts of each taxon to a simple presence/absence scheme. This resulted in nearly equivalent RF performance relative to training RF with the original continuous OTU inputs: ROC AUC of 0.862 (binary) vs 0.882 (continuous) and PR AUC of 0.846 (binary) vs 0.886 (continuous) (Figure 3A-B). This is an important result because it allows us to think about associations just in terms of presence or absence of an OTU without sacrificing much in model performance. We applied BowSaw to the Crohn’s disease samples and visualized 56,902 resultant sub-rules ranging in complexity from 2 to 7 variables (Figure 3C). There were 1,941 sub-rules with *F* = 1. We selected the most general of these rules (max|***I***|) to be the top candidate for the curate algorithm and found that it considers the status of 5 OTUs and accounts for 38 of 86 Crohn’s disease samples (Figure 3C). We set an association threshold of 90% and ended up with 10 sub-rules that together account for all 86 Crohn’s disease samples and an additional 11 non-Crohn’s disease samples (4 non-IBD, 7 ulcerative colitis). The top five rules combine to account for 78 of 86 Crohn’s disease samples and include 10 non-Crohn’s disease samples (Table 1).

**Table 1.** Association rules identified by BowSaw that account for all Crohn’s disease samples.

| Rule | CD Samples | Non CD Samples | New Samples Covered | Taxonomy | Presence |
| --- | --- | --- | --- | --- | --- |
| 1 | 38 | 0 | 38 | *Bacteroides* (genus) | y |
|  |  |  |  | *Lachnolostidium* (genus) | y |
|  |  |  |  | *Tyzzerella* (genus) | n |
|  |  |  |  | *Lachnospira* (genus) | n |
|  |  |  |  | *Lachnospiraceae* UCG-001 (genus) | n |
| 2 | 41 | 4 | 20 | *Dialister* (genus) | y |
|  |  |  |  | *Christensenellaceae* R7 group (genus) | n |
|  |  |  |  | *Christensenellaceae* R7 group (genus) | n |
|  |  |  |  | *Collinsella* (genus) | n |
|  |  |  |  | *Ruminococcaceae* (family) | n |
|  |  |  |  | *Finegoldia* (genus) | n |
|  |  |  |  | *Ruminococcus 1* (genus) | n |
| 3 | 9 | 1 | 9 | *Ruminococcus 1* (genus) | y |
|  |  |  |  | *Ruminococcaceae* UCG-002 (genus) | n |
|  |  |  |  | *Lachnospiraceae* (family) | n |
| 4 | 24 | 2 | 6 | *Streptococcus* (genus) | y |
|  |  |  |  | *Tyzzerella* (genus) | n |
|  |  |  |  | *Lachnospiraceae* (family) | n |
|  |  |  |  | *Hafnia* *Obesumbacterium* | n |
| 5 | 27 | 3 | 5 | *Lachnospiraceae* UCG-008 (family) | y |
|  |  |  |  | *Ruminococcus 1* (genus) | n |
|  |  |  |  | *Eubacterium eligens* group | n |
| 6 | 5 | 0 | 2 | *Ruminococcus 1* (genus) | y |
|  |  |  |  | *Dorea* (genus) | n |
| 7 | 7 | 0 | 2 | *Bacteroides* (genus) | y |
|  |  |  |  | *Dialister* (genus) | n |
|  |  |  |  | *Eubacterium rectale* group | n |
| 8 | 15 | 0 | 2 | *Lachnospiraceae* NK4A136 group | y |
|  |  |  |  | *Eubacterium eligens* group | y |
|  |  |  |  | *Tyzzerella* (genus) | n |
|  |  |  |  | *Christensenellaceae* R7 group (genus) | n |
|  |  |  |  | *Lachnospira* (genus) | n |
| 9 | 3 | 0 | 1 | *Ruminococcus gnavus* group | y |
|  |  |  |  | *Veillonella* (genus) | n |
|  |  |  |  | *Bacteroides* (genus) | n |
|  |  |  |  | *Finegoldia* (genus) | n |
| 10 | 10 | 1 | 1 | *Parabacteroides* (genus) | y |
|  |  |  |  | *Eubacterium eligens* group | y |
|  |  |  |  | *Ruminococcaceae* UCG-003 (genus) | n |
|  |  |  |  | *Lachnospiraceae* ND3007 group | n |

The top candidate rule is comprised of the presence of *Bacteroides* and *Lachnoclostridium* and the absence of three genera from the family *Lachnospiraceae: Lachnospira*, *Tyzerrella*, and *Lachnospiracea UCG 001* (Figure 3D). Detection of *Bacteroides* was nearly ubiquitous within the cohort, it was found in 170 of 178 total samples, but only 3 of the samples in which it was missing are diagnosed as Crohn’s disease. For the remaining taxa we performed a t-test comparing the distribution of the taxa in Crohn’s disease versus ulcerative colitis and versus healthy samples. *Lachnoclostridium* was frequently found in Crohn’s disease (67/86) but not in ulcerative colitis (27/46, p = .02) and was detected at roughly the same rate in non-IBD samples (34/46, p = .616). Detection of *Lachnospira* was depleted in Crohn’s disease samples (20/86) relative to ulcerative colitis (20/46, p = .022) and to non-IBD samples (31/46, p = 9.9^−7^). *Tyzzerella* was also detected at a lower rate in Crohn’s disease (63/86) relative to ulcerative colitis (24/46, p = .019) and non-IBD (24/46, p = .019). *Lachnospiracea UCG 001* was rarely detected in Crohn’s disease (4/86) which is a lower rate than it was detected in ulcerative colitis (9/46, p = .022) and in non-IBD samples (19/46, p = 1.45^−5^).

## Discussion

Interpretation of random forest models for classification may be confounded when there are multiple rules (combinations of variables and their specific values) associated with a phenotype of interest. We have developed BowSaw, which is an algorithmic approach for identifying the rules that a trained random forest model uses to make classifications when the values are categorical in nature. By taking advantage of the structure of trees found within a random forest, BowSaw produces a set of multiple decision rules that combine to account for each sample with a given observed phenotype. When the variables are the presumed causal agents, these rules represent plausible mechanistic relationships.

Results on simulated data demonstrate that when there are multiple rules associated with a single phenotype label that BowSaw is capable of faithfully identifying them. Application to data from the human microbiome project offers further evidence that BowSaw provides an efficient way of generating plausible hypotheses for high through put metagenomics studies. In particular we identified a rule that utilizes a presence/absence pattern of five microbial taxa (present: *bacteroides*, *lachnoclostridium*, absent: *lachnospira*,*lachnospiracea*, *tyzerrella*) that accounts for nearly half of all Crohn’s disease samples in the cohort (38/86). This specific pattern of microbial colonization in the guts of Crohn’s disease patients is unreported, but each taxon’s respective enrichment or depletion status and association with disease status has been reported. If the cohort of patients in the human microbiome study are representative of all people afflicted by Crohn’s disease then this rule represents a significantly large sub-set of those suffering. Inquiries into the relationship of the taxa included in this rule with disease status may yield important insights into the mechanisms of the disease and potential therapeutic strategies for this sub-population. Of the five associated taxa, we suspect that the absence of *lachnospira*, *lachnospiracea UCG 001*, and *tyzzerella* are biologically meaningful. We have reason to believe so because it has been reported that the *lachnospiraceae* family is generally suppressed in Crohn’s disease [32–34]. *Lachnospira* has been reported as depleted with respect to Crohn’s disease several times [35, 36]. The depletion of *tyzzerella* has been associated with chronic intestinal inflammation and supplementation suggested as a probiotic for Crohn’s disease [37, 38]. While the relationship of *lachnospiracea UCG 001* with Crohn’s disease is still unclear, its depletion has been reported in mice displaying symptoms of anhedonia and it was significantly enriched in anhedonia resilient mice [39]. Partly because IBD is frequently accompanied by depression, anhedonia has been suggested as an important symptom in the diagnosis of IBD [40]. The associations of the individual OTUs defined by this rule are consistent with previously reported findings in the existing literature and describe a taxonomic profile that exclusively identifies a large sub-population of Crohn’s disease samples within this cohort. The presence of *bacteroides* does not appear to be particularly useful and in this context is probably preserved because it causes a perfect association, although high levels of some species are implicated in the pathology of Crohn’s disease [41]. *Lachnoclostridium*, is differentially distributed across the three classes. Notably it is less frequently detected in ulcerative colitis relative to Crohn’s and non-IBD samples, which roughly resemble one another. Increased levels of this genus was detected in rats that showed relief of colitis symptoms after treatment with a proposed therapeutic agent [42].

The current implementation of the algorithms are restricted to classification tasks with categorical predictor values, this is a challenge that we will need to address in order to make the approach more generally applicable. Future work will also focus on extending these for the interpretation of regression models. Such additions will greatly increase the number of systems to which we can apply BowSaw.

## Acknowledgments

We are grateful to members of the Segrè lab for helpful discussions and for feedback on the manuscript. DS and DD acknowledge funding from the Defense Advanced Research Projects Agency (Purchase Request No. HR0011515303, Contract No. HR0011-15-C-0091), the U.S. Department of Energy (DE-SC0012627), the NIH (T32GM100842, 5R01DE024468, R01GM121950 and Sub_P30DK036836_P&F), the National Science Foundation (1457695), the Human Frontiers Science Program (RGP0020/2016), and the Boston University Interdisciplinary Biomedical Research Office. DD is grateful to Dr. Nisha Rajagopal for her patience in conversations about random forests and her valuable insight.

