## Supplemental Figure 1 for "BowSaw: inferring higher-order trait interactions associated with complex biological phenotypes"

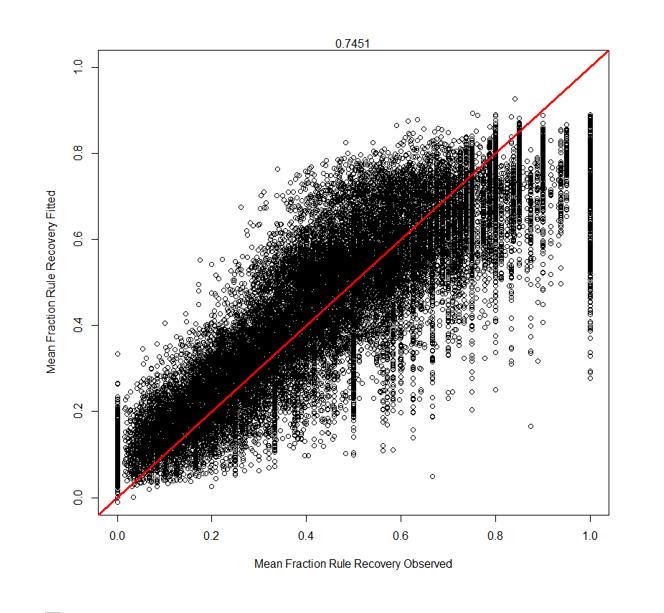

Supplemental Figure 1 There is a strong linear relationship between the performance of BowSaw and observable model evaluation metrics measured here as the average fraction of a complete rule recovered in any candidate rule (y). The linear model specified was: y = sampleSize + #features + ROCAUC + PRAUC.
